# Lempel-Ziv complexity of the EEG predicts long-term functional recovery after stroke in rats

**DOI:** 10.1101/248039

**Authors:** Susan Leemburg, Claudio L. Bassetti

**Affiliations:** Department of Neurology, University Hospital Zurich, Zurich, Switzerland; Neurology Department, University Hospital Bern, Switzerland; Medical Faculty, University of Bern, Switzerland

## Abstract

Non-linear complexity of the EEG signal can be used to detect abnormal brain function relating to behavioral deficits. Here, we compare the effects of experimental stroke on EEG complexity using Lempel-Ziv complexity analysis (LZC) and multiscale entropy analysis (SampEn).

EEG was recorded in bilateral motor cortex at baseline and during a 30-day recovery period after distal middle cerebral artery occlusion in rats. Motor function was assessed using a single pellet reaching task. Stroke caused an acute drop in both LZC and SampEn in the ipsilesional hemisphere in wakefulness, NREM and REM sleep, as well as reduced pellet reaching success. SampEn reductions persisted for at least 10 days post-stroke, whereas LZC had returned to baseline levels by day 4. EEG complexity in the contralesional hemisphere and in sham-operated animals were unaffected.

If EEG complexity reflects post-stroke brain function, post-stroke asymmetry could be used to predict behavioral recovery. In rats, acute LZC asymmetry was significantly correlated with the amount of motor function recovery by post-stroke day 31, but SampEn asymmetry was not. EEG LZC may thus be a useful tool for predicting functional recovery after stroke. MSE could be effective in identifying cortical dysfunction, but does not reflect behavioral outcomes.

## Introduction

Stroke remains a common cause of long-term disability in industrialized countries [1]. Predictive models can aid in the planning and implementation of rehabilitation strategies and help optimize functional recovery [2,3]. EEG, as an accessible and noninvasive method of measuring brain function, could be used to complement behavioral post-stroke assessments used to guide rehabilitation strategies.

Post-stroke effects on EEG power and coherence spectra have been described previously, but their relation to functional recovery is not straightforward [4–6]. The use of spectral power to assess brain function after stroke is further complicated by inter individual differences. Moreover, both overall spectral power and power within specific frequency bands vary with age and sex and are affected by pharmacological treatment and sleep-wake history [7–9]. Linking EEG power spectra to behavioral function after stroke is further complicated by large inter-individual differences in power spectra and limited information about the healthy EEG of most patients.

Non-linear complexity measures, such as Lempel-Ziv complexity (LZC) [10] and multiscale entropy (MSE) [11,12] yield information about the EEG signal that is independent of amplitude. Nonlinear complexity measures can be more sensitive than spectral power in detecting abnormal brain function in epilepsy [13], Alzheimer’s disease [14–17], depression, schizophrenia [18,19] and after brain injury [20], as well as sleep homeostasis in healthy rats [21]. Moreover, post-stroke LZC was a more accurate predictor of post-stroke depression than spectral power in patients [22]. Similarly, LZC predicted behavioral responsiveness based on brain function in anaesthetized patients with greater accuracy than approximate entropy, spectral entropy, or median EEG frequency [23]. MSE can be used to detect abnormal brain activity in damaged cortical areas in chronic stroke [24]. Additionally, MSE might be used to predict recovery in a rat model of cardiac arrest [25].

Stroke-related changes in EEG complexity have been described in stroke patients. Hemispheric stroke led to reduced EEG complexity in the ipsilesional hemisphere in stroke patients [26], but was increased after thalamic stroke [27]. EEG complexity calculated was thus differentially affected by lesion location, but also by the methods used to compute it [22,24,26,27]. Here, we compare how distal middle cerebral artery occlusion affects EEG complexity calculated using LZC or MSE during a 30-day recovery period after in rats, as well as the use of these measures in predicting motor function recovery.

## Materials and Methods

### Experimental setup

15 rats were implanted with EEG and EMG electrodes after 3 weeks of training on the SPR task. After electrode implantation, they were allowed to recover for 8-9 days, during which training continued. 24-hour baseline EEG recordings were then performed, as well as baseline skilled reaching (SPR) tests. Focal cortical ischemia by middle cerebral artery occlusion (MCAO) was then induced in the hemisphere contralateral to the rat’s preferred paw in 8 animals. 7 Sham operated animals served as a control group. Post-surgery EEG recordings were performed on day 1, 4, 7, 10, and 30. The first post-surgery EEG recording (D1) was performed approximately 20h after surgery. Post-surgery SPR testing was performed on days 2, 5, 8, 11, and 31. Rats were sacrificed after the final SPR session. Mean LZC and SampEn were calculated for wake, NREM and REM for each animal to assess effects of stroke on temporal complexity of the EEG signal.

### Animals

15 male Sprague-Dawley rats were used (180-200 g at the start of the experiment, 8 MCAO, 7 sham, Harlan). Rats lived in individual plexiglass cages on a 12h:12 h light-dark cycle. They were fed 20-25g of chow once per day at lights-on, or after completing motor skill testing or training (SPR). The feeding schedule did not result in significant weight loss. On SPR testing or training days, rats received chocolate flavored dustless precision pellets (45 mg, #F0299, Bioserv Inc.) in addition to the regular chow. The number of pellets a rat consumed depended on SPR success, but did not exceed 50. Water was available *ad libitum* throughout the experiment. All experiments were carried out in the University Hospital Zurich, Switzerland, according to local regulations for the care and use of laboratory animals and with governmental approval (Licenses 167/2005 and 190/2008, Kantonales Veterinäramt Zürich, Switzerland).

### Assessment of motor function

Fine motor function was assessed using a skilled reaching task (SPR), in which a rat retrieved a small food pellet located outside of the testing cage by reaching for it using one forepaw, as described previously [28,29]. The SPR testing cage consisted of a rectangular plexiglass box with small shelves mounted 3 cm above the cage floor on the outside of each short wall, which could be reached by the rats via a 1.4 cm wide window in the cage wall. Small indentations in the shelves aligned with the edges of the window assured constant placement of pellets on the shelves (1.5 cm from the inside of the cage).

The first week of three weeks of SPR training consisted of daily 10-minute sessions, in which pellets initially placed within easy reach of the rat. Once the rat reliably consumed these pellets, they were moved progressively farther away to encourage reaching with the forepaw. Once a rat demonstrated paw-preference by making more than half of the reaching attempts in a session with a single paw, pellets were placed in the indentation that could only be reached with that preferred paw. In week 2 and 3 of training, in which pellets were presented on alternating sides of the reaching cage. As a result, the rat fully repositioned its body for each reaching attempt. Training sessions ended after the rat had performed 50 reaching attempts, or after 15 minutes.

During baseline and post-MCAO sessions, pellets were only presented on one side of the cage after the rat had turned away from the window and moved to the other side of the cage. SPR attempts were classified as either successful (the rat grasped the pellet, transferred it into its mouth and ate it) or failed (the rat dropped the pellet or knocked it off the shelf). Testing sessions were videotaped for later verification. SPR success was calculated as the percentage of successfully obtained pellets out of 50 possible attempts.

### Electrode implantation and MCAO surgery

During all surgical procedures, rats were anaesthetized using 2-2.5% isoflurane in 30% oxygen and 70% N_2_O and rectal temperature was maintained at 36°C.

EEG electrodes (gold-plated miniature screws) were implanted epidurally in each hemisphere (rostral electrode: bregma +2 mm, 2 mm lateral; caudal electrode: bregma −2 mm, 2.5 mm lateral). The caudal electrode served as reference for the rostral one in each hemisphere. Thus, the recorded area covered the intact motor cortex, but not the infract in the somatosensory cortex (Fig. 3A). Two gold wires inserted in the neck muscles served as EMG electrodes.

Ischemic stroke was induced by permanent occlusion of the distal branch of the middle cerebral artery (MCA), as described previously [30]. In short, the skull overlying the MCA was exposed through a vertical incision in the temporal muscle. A 4-by-5mm craniotomy was then made in the frontal bone, exposing the brain and middle cerebral branches. After carefully opening the dura mater, the MCA and its main branches were closed using bipolar electrocoagulation. Finally, the temporal muscle and overlying skin were sutured back in place. Sham operated rats underwent the same procedure, but without opening the dura mater and coagulation of the MCA. MCAO and sham surgeries were performed in the hemisphere contralateral to the rats’ preferred reaching forepaw.

### EEG recording

EEG and EMG activity were recorded using an Embla A10 amplifier and Somnologica Science software (Embla) at 100 Hz and 200 Hz sampling rates, respectively. The EEG signal was filtered using a low-cut filter at 0.3 Hz. EMG was filtered for 50 Hz artefact, and had a low-cut filter at 10 Hz.

EEG was scored as wake, NREM or REM in 8-second epochs based on the contralesional EEG signal and on the EMG using Somnologica Science (Embla). Wherever the contralesional EEG contained many artefacts, the signal from the ipsilesional hemisphere was consulted to help identify the vigilance state. Throughout the experiment, wake was characterized by a low amplitude, high frequency EEG pattern and high EMG activity. NREM was characterized by the occurrence of high amplitude slow waves and tonic low EMG activity. The low amplitude EEG activity during REM was dominated by theta activity and only occasional twitches were present in the EMG signal. Epochs were classified as belonging to a vigilance state when more than half of the epoch matched the criteria for that state. 22 hours of each recording day were analyzed, beginning 1 hour after the start of the recording at lights-on. Only artifact-free epochs were included in analyses.

### Data analysis

#### Lempel-Ziv complexity

Lempel-Ziv complexity (LZC) is a nonparametric measure of temporal complexity of the EEG signal. This method estimates the complexity of a finite series of numbers by computing the number of distinct subsequences within that series (Fig. 1A). LZC was calculated based on the methods described by Lempel and Ziv (1976) [10]. First, each artifact-free EEG epoch was converted into a binary sequence by coarse-graining the signal based on the median signal value for the epoch [23]. Each data point *x*(*n*) of the original 8-second EEG epoch was compared to epoch median *M*, resulting in a new binary 800-point series *s* as follows: 
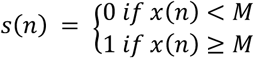
. The median was chosen for coarse-graining because of its robustness in case of outliers. This resulting binary series was then scanned from left to right and a complexity counter *c*(*n*) was increased whenever a new subsequence of consecutive characters was detected (for a detailed description see Zhang et al. (2001), [23]). As complexity *c*(*n*) depends on both epoch length *n*, (here, *n* = 800) and the number of different symbols within the coarse-grained series, *α* (here *α* = 2 for a binary series), *c*(*n*) was normalized using the theoretical upper bound *b* for *c*(*n*), calculated as 
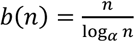
. A normalized complexity value that is independent of epoch length was then calculated as 
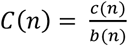
. This normalized value is used whenever the Lempel-Ziv complexity (LZC) of a signal is discussed in the current paper. Mean LZC of all artifact-free epochs on a recording day was then calculated separately for wake, NREM and REM for every rat.

**Figure 1:**
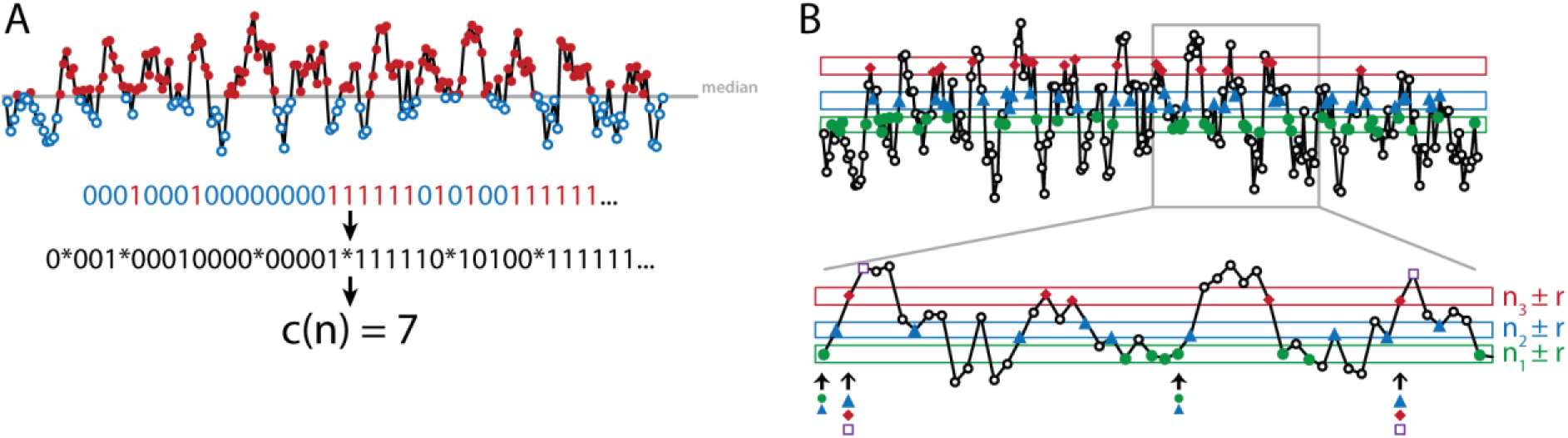
Comparison of Lempel-Ziv and multiscale entropy analysis. **A:** Lempel-Ziv complexity analysis. Signals are coarse-grained to form a binary sequence by classifying each data point as ≤ median (open circles, blue) or > median signal amplitude (closed circles, red). Unique subsequences are then detected as described [10]. Lempel-Ziv complexity of the sequence is defined as the number of detected unique subsequences c(n). **B:** Multiscale entropy analysis. MSE is used to calculate the probability of a template sequence of two data points repeating within the analyzed signal (m = 2). To this end, data points are classified as being equal within a tolerance r (i.e. 20% of the signal standard deviation). Here, this is shown for data point n_1_ (closed circle, green), n_2_ (triangle, blue) and n_3_ (closed diamond, red); boxes around these data points indicate n_1_ ± r, n_2_ ± r and n_3_ ± r. Two data points are considered equal if the absolute difference between them is < r. As such, data points matching n_1_ are shown as green circles. Points that match n_2_ and n_3_ are shown as blue triangles and red diamonds, respectively. To calculate Sample entropy (SampEn) of this signal, data points n_1_ and n_2_ will compose the first 2-datapoint template sequence n_1_n_2_. For the signal shown, this sequence is present once more. The matching 3-datapoint template sequence n_1_n_2_n_3_ is not detected in this example. This procedure is repeated for each next pair of 2-datapoint and 3-datapoint template sequences (n_2_n_3_ and n_2_n_3_n_4_, n_3_n_4_ and n_3_n_4_n_5_, etc.). In this example, n_2_n_3_ and n_2_n_3_n_4_ are each detected twice. The number of sequences that match each of the 2‐ and 3‐ data point template sequences are added to the previously counted matches. Thus, for the first 4 data points in this signal, the number of detected 2-datapoint sequences is 4 and the number of detected 3-datapoint sequences is 2. This procedure is repeated for each pair of 2-datapoint and 3-datapoint template sequences in the signal. SampEn is calculated as the natural logarithm of the ratio between the number of detected 2‐ and 3-datapoint sequences and reflects the probability that sequences that match each other for the first two data points will also match for the next point.

#### Multiscale Entropy

Like LZC, multiscale entropy analysis is used to assess the temporal complexity of the EEG signal. Unlike LZC, for which the number of unique sequences within a series are counted, multiscale entropy analysis assesses the complexity of a signal by estimating the likelihood of repeating sequences occurring within the signal (Fig. 1B), reflected by the signal's sample entropy (SampEn) [11,12]. The calculation of SampEn depends on parameters *n*, *m*, *τ,* and *r*. *n* denotes the number of data points in the analyzed series (here, *n* = 800 for an 8-second EEG epoch). *m* defines the length of the repeating sequences. Here we used *m = 2*. That is, we calculated the likelihood of 2 subsequent values (a template sequence) occurring repeatedly within the epoch. This was repeated for each pair of values in the epoch. *r* and *τ* are coarse-graining parameters. *τ* specifies coarse-graining within the time-dimension of the series; here we analyzed the original EEG signal using *τ= 1*. *r* is a tolerance value controlling the level of similarity between data points in the amplitude-dimension of the EEG signal. Data points are considered equal if the absolute difference between them was less than or equal to *r*. In this paper, *r* was set as 20% of epoch standard deviation.

SampEn is calculated as follows:

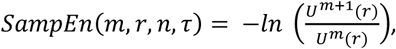

or the likelihood of an m-length sequence (for *m = 2*, 2 data points) repeating relative to the likelihood of a repeating m+1-length sequence (for *m = 2*, 3 data points).

*U^m^*(*r*) is calculated as: 
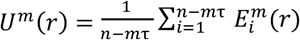
.
*U^m^*(*r*) is based on probability function 
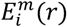
, formed by the number of m-length sequences 
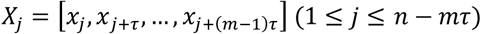
 in an epoch that are considered template sequence *X_i_*(*i* ≠ *j*), so that 
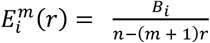
.

*B_i_* is equal to the number of sequences *X_j
_* whose difference from template *X*_*i*_ is less than or equal to *r*. That is, *B_i_* is equal to the number of sequences that were equal to the template sequence. After SampEn was calculated for each artifact-free EEG epoch, mean SampEn of all artifact-free epochs on a recording day was calculated separately for each vigilance state for every rat.

#### Relation between complexity and power spectra

Power spectra for frequencies from 0.5 to 25 Hz were calculated for artifact-free epochs in wakefulness, NREM and REM in baseline EEG recordings (N = 15 animals; 8 MCAO and 7 sham) using Welch’s power spectral density estimate with Hamming window and 50% overlap between epochs (pwelch in Matlab 2015b). Pearson's correlation analysis was then performed between mean LZC or SampEn in a vigilance state and spectral power for each frequency bin. Correlations were considered significant when p < 0.05.

#### Simulated signals

Relations between LZC and SampEn were assessed in wake EEG signals, as well as in three different simulated signals generated in Matlab 2015b: a 15Hz sine wave sampled at 100 Hz, Gaussian white noise and 1/f noise. The latter two were generated using the the wgn function and dsp.ColoredNoise, respectively. LZC and SampEn were calculated for 100 epochs of 800 data points for each simulated signal.

#### Statistical Analyses

Time and treatment effects on SPR performance, LZC and SampEn were analyzed using repeated measures ANCOVA. Baseline values were included as covariates to control for possible confounding effects of baseline differences. Greenhouse-Geisser sphericity corrections were applied where appropriate. Correlations were calculated using Pearson's R. Effects were considered significant when p < 0.05. Statistical analyses were performed in JASP v0.8.0.1 and Matlab 2015b. Values are presented as mean ± s.e.m. unless stated otherwise.

### Histology

Rats were killed with an overdose of sodium pentobarbital and perfused transcardially with 0.1M phosphate-buffered saline (PBS) followed by 4% paraformaldehyde (PFA) in 0.1M PBS. Brains were removed, post-fixated in 4% PFA in 0.1M PBS and cryoprotected in 30% sucrose in 0.1M PBS at 4°C. 30μm thick coronal sections were collected and stained using cresyl violet. Every 12th section was stained and analyzed. Chemicals were purchased from Sigma-Aldrich. Histological results were analyzed using ImageJ software (U.S. National Institutes of Health). Infarcts were outlined based on the absence of Nissl-stained neurons in the cortex. Infarct volume was calculated as follows:

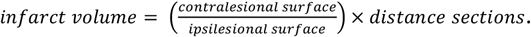

## Results

### Relation between LZC, MSE and power spectrum

Although Lempel-Ziv complexity analysis and multiscale entropy analysis are both used to assess the temporal complexity of a signal, they differ in their approach (Fig. 1). As a result, LZC and SampEn of a signal are not correlated in a straightforward manner. To investigate the relation between the two, we compared LZC and SampEn in four signal types with different regularity and complexity (sine, white noise, 1/f noise and wake EEG; Fig. 2A).

**Figure 2:**
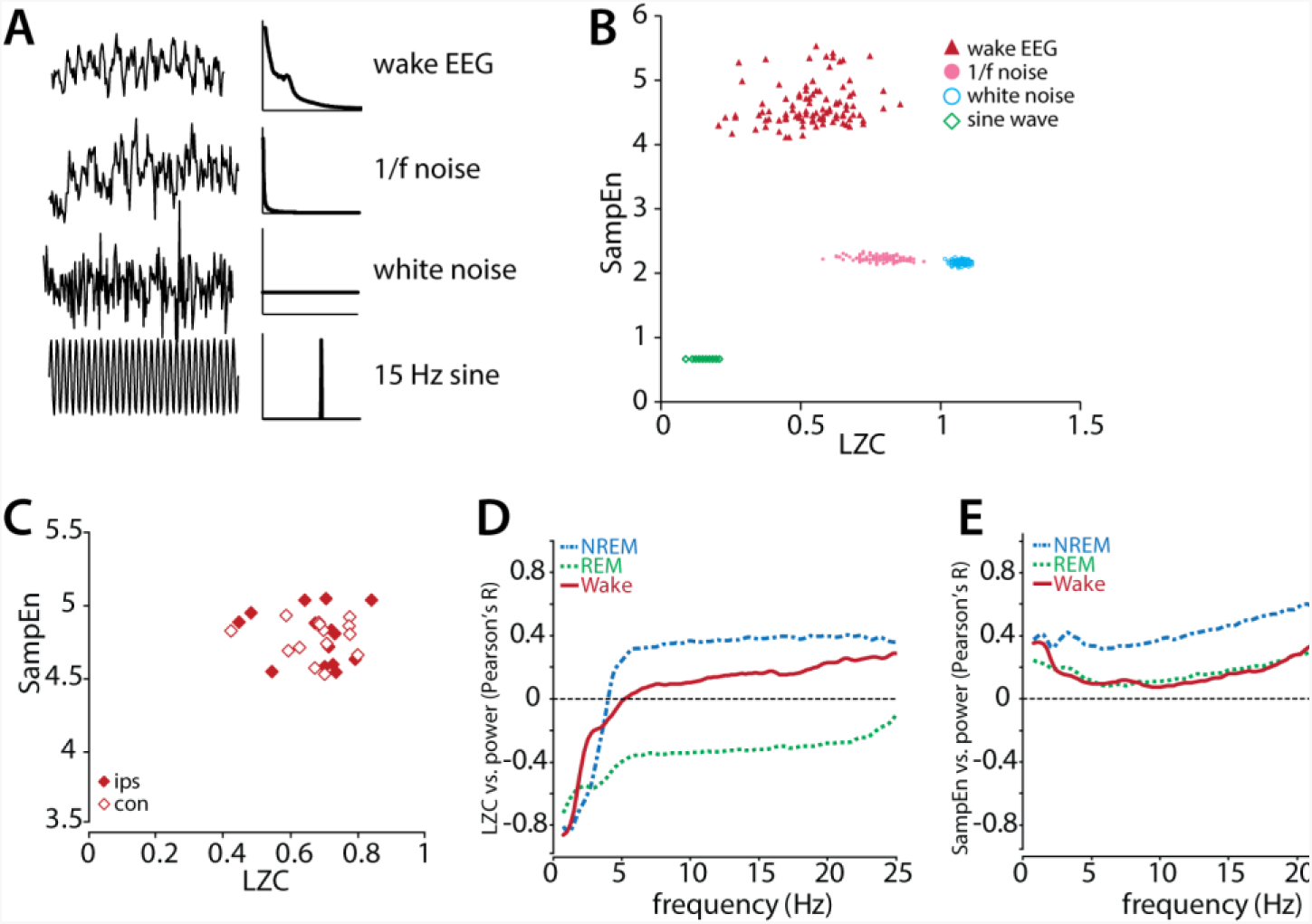
Relation between LZC and MSE in simulated and real signals. **A:** Comparison of waveforms and power spectra of wake EEG and simulated signals (1/f noise, white noise and sine wave). **B:** Relation between LZC and SampEn in simulated signals and wake EEG. Shown are 100 epochs for each signal type. All EEG epochs shown are from the same recording session in a representative animal. **C:** Relation between LZC and SampEn of wake EEG in trained (ips) and untrained (con) hemisphere at baseline. Shown are mean values for each animal (N = 15). **D:** Correlation between LZC and spectral power in baseline wake, NREM and REM EEG (N=15 rats). **E:** Correlation between SampEn and spectral power in baseline wake, NREM and REM EEG (N=15 rats).

As a simple and highly regular signal, the sine wave signal had very low LZC and SampEn (0.08 ± 0.002 and 0.681 ± 2.23*10^−17^, respectively). By contrast, white noise had high LZC (1.05 ± 0.001) and moderately high SampEn values (2.19 ± 0.003), being highly irregular, but by definition not containing any non-random structure within the signal. 1/f noise, an irregular signal that is defined by its frequency-amplitude relation, had lower LZC values than white noise (0.75 ± 0.005), but higher SampEn (2.23 ± 0.004). Finally, LZC of wake EEG (0.538 ± 0.013) in a representative animal was similar to that of 1/f noise, but its SampEn was much higher (4.615 ± 0.033). Within each of these four signals, LZC was not correlated to SampEn. Similarly, mean LZC and mean SampEn for each animal were not significantly correlated in the healthy rat brain (Fig. 2C, N = 15 animals, ips: R = −0.103, p = 0.727, con: R = −0.046, p = 0.876).

Thus, a signal can have high LZC, but low SampEn and vice versa. This is a result of the different methods by which LZC and MSE analyses assess temporal structure of the signals. Where LZC is mostly reflects the regularity of a signal, SampEn also reflects structure within a signal. Although EEG and 1/f noise were less complex than white noise when assessed by LZC, their SampEn is higher due to the presence of frequency-amplitude dependencies within these signals.

LZC and SampEn are not simply reflections of the low-or high-frequency components of a complex signal like the EEG. LZC was not significantly correlated to spectral power in the wake, NREM or REM (Fig. 2D). The relation between LZC and power varies with frequency: low frequency power (< 3 Hz) was negatively correlated to LZC and a weaker positive correlation in was present in frequencies 5-25 Hz in wake and NREM, although none of these were statistically significant (Fig. 2D). LZC-power correlations in REM sleep followed a similar pattern of non-significant stronger correlation in low frequencies and weaker correlations in higher frequencies.

Similar to LZC, SampEn was not significantly correlated to power in all analyzed frequencies in NREM, REM and wakefulness. In all three vigilance states, SampEn showed a weak positive correlation with power. Correlations were stronger in higher frequencies (20-25 Hz), but did not reach statistical significance (Fig. 2E).

### Stroke-related changes in histology and motor function

Middle cerebral artery occlusion (MCAO) resulted in infarcts located in the somatosensory cortex (Fig. 3A). Ischemic damage did not extend into subcortical regions, nor into cortical areas located directly adjacent to or below the EEG electrodes. Infarct volume was 23.9 ± 3.1 mm^3^ (N = 8, Fig. 3A).

**Figure 3:**
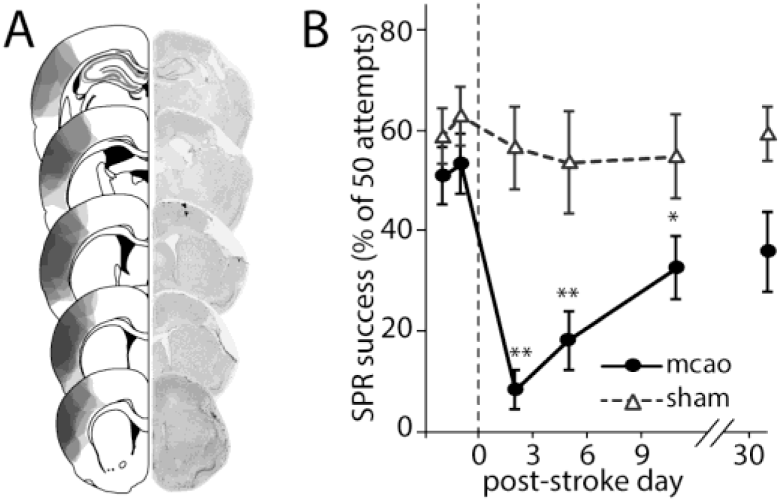
Histological and behavioral results. **A:** Location and extent of ischemic lesions after distal middle cerebral artery occlusion (MCAO). Outlines show lesions for individual animals (N = 8, left), photomicrographs (right) show the infarct of a representative animal. **B:** Motor function in MCAO and sham-operated animals pre-and post-surgery. Values are mean ± s.e.m, asterisks indicate significant difference between sham and MCAO (* p<0.05, **p<0.01).

MCAO led to a marked acute reduction in pellet reaching success (Fig. 3B; repeated measures ANCOVA, group effect: F(1) = 26.63, p < 0.001; time effect: F(2.98) = 10.65, p < 0.001; time*group interaction F(2.98) = 8.69, p < 0.001). Performance improved from day 2 through 11 and returned to baseline levels on D31. Sham surgery had no effect on SPR success.

### Effects of stroke on Lempel-Ziv complexity and sample entropy

Stroke acutely reduced LZC in ipsilesional EEG in MCAO rats in wakefulness (Fig. 4A; repeated measures ANOVA; group effect F(3) = 3.081, p = 0.046; time effect: F(4) = 6.939, p < 0.001; group*time interaction F(12) = 1.661, p = 0.087). Contralesional EEG in MCAO rats showed a small, non-significant reduction in LZC. Ipsilesional wake LZC had returned to sham and baseline levels by day 4.

**Figure 4:**
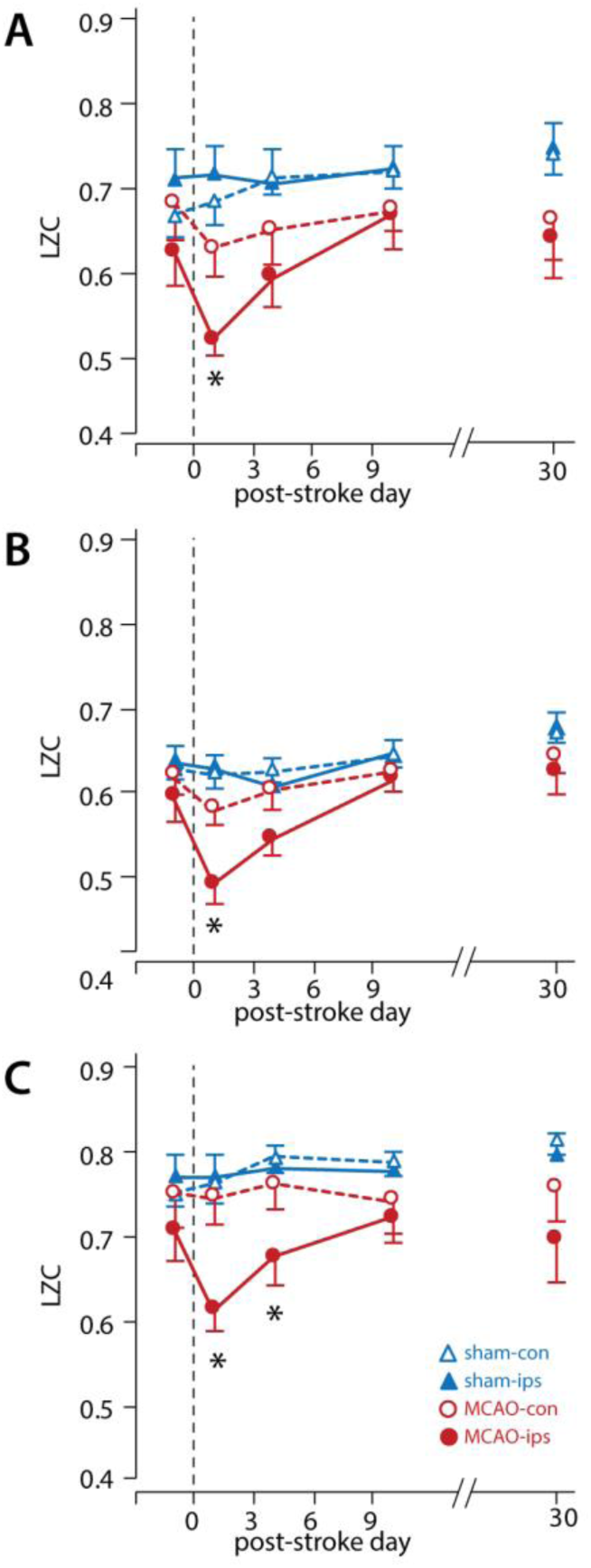
Effects of stroke on LZC. Time course of LZC during wake **(A)**, NREM **(B)**, and REM **(C)** in ipsilesional (ips) and contralesional (con) hemispheres of sham and MCAO animals. Values are mean ± s.e.m. Asterisks indicate significantly reduced values in MCAO-ips (*p<0.05)

Effects of stroke on NREM were similar to those found during wake (Fig. 4B; repeated measures ANOVA; group effect F(3) = 3.371, p = 0.034; time effect: F(2.943) = 7.796, p < 0.001; group*time interaction F(8.830) = 2.132, p = 0.038). NREM LZC returned to baseline levels by day 4 as well.

By contrast, ipsilesional reductions in REM lasted slightly longer and had recovered by day 10 (Fig. 4C; repeated measures ANOVA; group effect: F(3) = 3.077, p = 0.046; time effect: F(2.591) = 6.679, p < 0.001; group*time interaction F(7.774) = 1.535, p = 0.108).

Like LZC, SampEn was significantly reduced in the ipsilesional hemisphere of MCAO rats. This reduction was longer lasting than the stroke-related changes in LZC. Ipsilesional SampEn in wake was significantly reduced on day 1, 4 and 10, but returned to baseline levels by day 30 (Fig. 5A; repeated measures ANOVA; group effect F(3) = 9.648, p < 0.001; time effect: F(2.708) = 1.841, p = 0.127; group*time interaction F(8.125) = 5.795, p < 0.001). Effects of stroke on ipsilesional SampEn in NREM were similar to those found in wakefulness (Fig. 5B; repeated measures ANOVA; group effect F(3) = 9.746, p < 0.001; time effect: F(2.702) = 1.810, p = 0.159; group*time interaction F(8.105) = 5.427, p < 0.001). SampEn in REM was also significantly reduced ipsilesionally in MCAO animals (Fig. 5C; repeated measures ANOVA; group effect F(3) = 9.746, p < 0.001; time effect: F(3.029) = 1.761, p = 0.161; group*time interaction F(9.087) = 5.790, p < 0.001).

**Figure 5:**
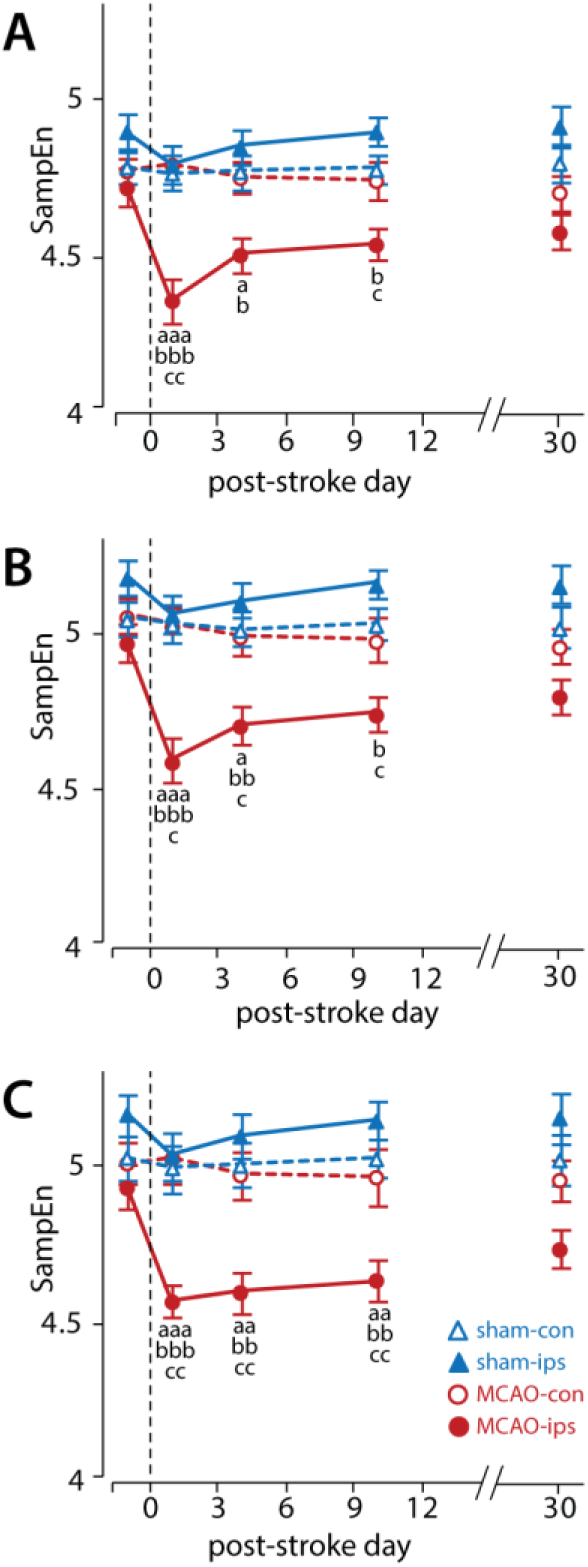
Effects of stroke on MSE. Time course of SampEn during wake **(A)**, NREM **(B)**, and REM **(c)** in ipsilesional (ips) and contralesional (con) hemispheres of sham and MCAO animals. Values are mean ± s.e.m. Letters denote significant differences between groups: MCAO-ips vs. MCAO-con: ^a^ p < 0.05, ^aa^ p < 0.01, ^aaa^ p ≤ 0.001, MCAO-ips vs. sham-con: : ^b^ p < 0.05, ^bb^ p < 0.01, ^bbb^ p ≤ 0.001, MCAO-ips vs. sham-ips: ^c^ p < 0.05, ^cc^ p < 0.01, ^ccc^ p ≤ 0.001.

### Relation between EEG complexity and motor function recovery

EEG complexity measures have previously been linked to altered brain function in stroke patients [22,24,26,27], although their relation to behavioral outcomes has not been studied. LZC and SampEn might be useful to help predict functional recovery after stroke (i.e. recovery of motor function). To this end, LZC and MSE asymmetry on day 1 post-stroke were correlated with the amount of SPR improvement from day 2 to 31. LZC and SampEn asymmetry were calculated as the ratio of contralesional to ipsilesional values for each animal, so that a large ipsilesional reduction resulted in a high asymmetry score.

Even though stroke-related changes in LZC were more transient than those found in SampEn, acute LZC asymmetry predicted the amount of functional recovery on D31 (Fig. 6A; wake R = −0.72, p = 0.021; NREM: R = −0.64, p = 0.048; REM: R = −0.71, p = 0.023). By contrast, MSE asymmetry had no relation to functional recovery (Fig. 6B; wake R = 0.11, p = 0.796; NREM: R = −0.19, p = 0.651; REM: R = 0.07, p = 0.874).

**Figure 6:**
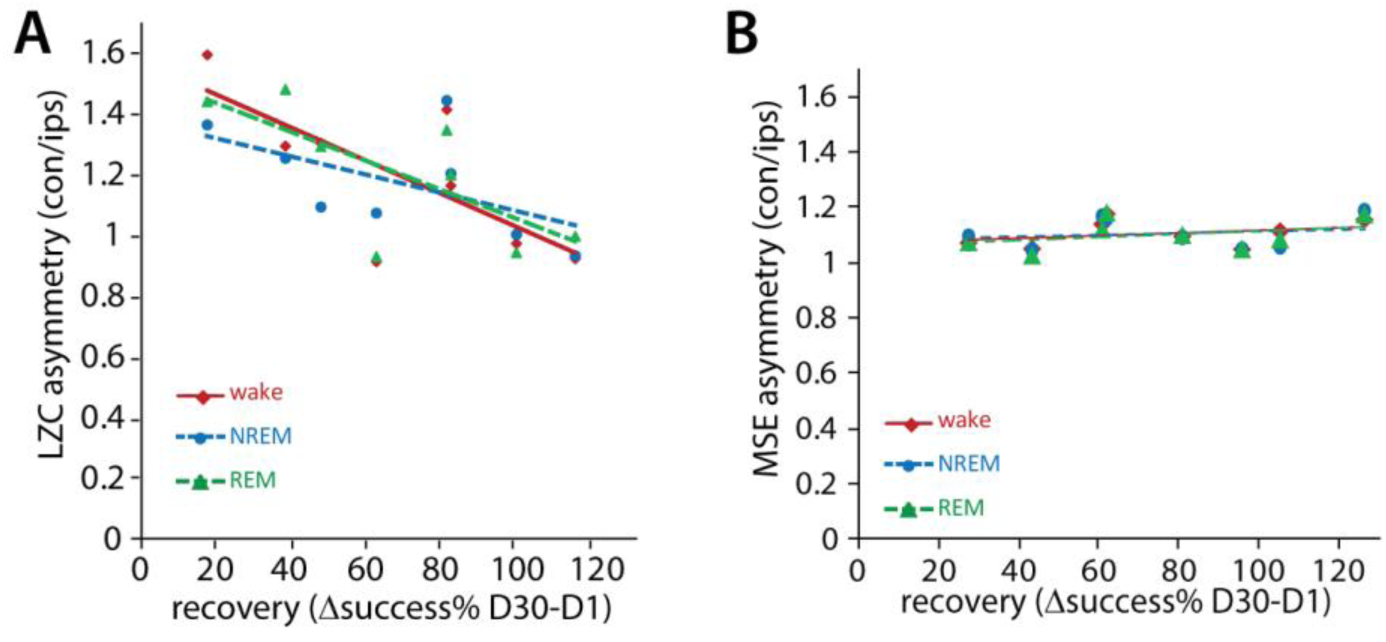
Correlation between functional recovery and EEG signal complexity. **A:** Correlation between acute LZC asymmetry (contralesional/ipsilesional on D1) and amount of motor function recovery on D30 (SPR success as % of baseline, ΔD30-D2) in wake (R = −0.72, p = 0.02), NREM (R = −0.64, p = 0.048) and REM (R = −0.71, p = 0.02). **B:** Correlation between acute SampEn asymmetry (MSE, contralesional/ipsilesional on D1) and degree of motor function recovery on D30 (SPR success as % of baseline, ΔD30-D2) in wake (R = 0.11, p = 0.79), NREM (R = −0.19, p = 0.65) and REM (R = 0.07, p = 0.87).

### Stability of LZC and MSE during the day

If LZC or SampEn are to be useful predictors of behavioral outcome after stroke, they ideally show little circadian and other non-stroke related variation. Wake LZC slightly increased during the first 4 hours of recording, although this increase was not statistically significant and variability was high (Fig. 7A; repeated measures ANOVA; group effect F(3) = 1.372, p = 0.275; time effect: F(2.205) = 22.344, p < 0.001; time*group interaction: F(6.614) = 1.326, p = 0.259). Likewise, no significant time effects were found in NREM in the first 4 hours of the day (Fig. 7B; repeated measures ANOVA; group effect F(3) = 1.426, p = 0.260; time effect: F(1.305) = 1.183, p = 0.300; time*group interaction: F(3.914) = 0.987, p = 0.105). LZC remained stable throughout the 22-hour recording period in both wake (Fig. 7C) and NREM (Fig. 7D) as well. This was the case in the healthy brain at baseline and one day post-stroke. Interhemispheric LZC asymmetry (contralesional/ipsilesional LZC) was not significantly affected by time of day at baseline or on day 1 post-stroke (Fig. 7D, E).

**Figure 7:**
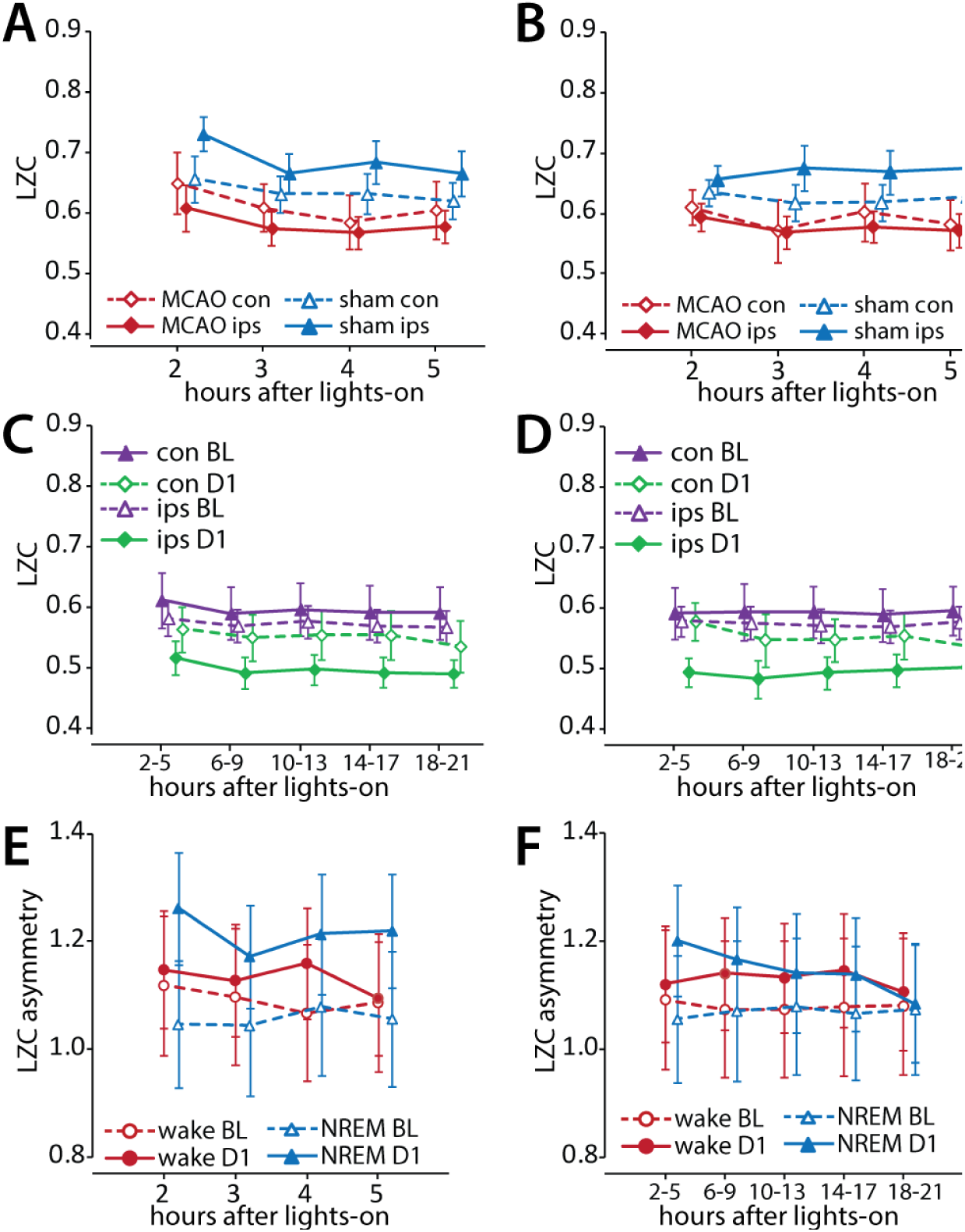
Circadian effects on LZC and LZC asymmetry. **A:** Mean wake LZC in the first 5 hours after lights-on in ipsilesional (ips) and contralesional (con) hemispheres of stroke (MCAO) and sham animals at baseline. Values are mean ± s.e.m. **B:** Mean NREM LZC in the first 5 hours after lights-on in ipsilesional (ips) and contralesional (con) hemispheres of stroke (MCAO) and sham animals at baseline. Values are mean ± s.e.m. **C:** Mean wake LZC in 4-hour bins at baseline (BL) and on D1 in MCAO animals. ips = ipsilesional, con = contralesional. Values are mean ± s.e.m. **D:** Mean wake LZC in 4-hour bins at baseline (BL) and on D1 in MCAO animals. ips = ipsilesional, con = contralesional. Values are mean ± s.e.m. **E:** MSE asymmetry in wake and NREM (contralesional/ipsilesional LZC) during the first 5 hours after lights-on in MCAO animals at baseline (BL) and on D1. **F:** MSE asymmetry in wake and NREM (contralesional/ipsilesional LZC) in 4-hour bins after lights-on in MCAO animals at baseline (BL) and on D1.

In contrast to LZC, SampEn in wake increased during the first 4 hours of recording (Fig. 8A; repeated measures ANOVA; group effect F(3) = 2.019, p = 0.138; time effect: F(3) = 29.296, p < 0.001; time*group interaction: F(9) = 1.028, p = 0.427). A concurrent decrease of similar magnitude was found in NREM sleep (Fig. 8B; repeated measures ANOVA; group effect F(3) = 1.316, p = 0.292; time effect: F(3) = 115.73, p < 0.001; time*group interaction: F(9) = 1.513, p = 0.160), possibly reflecting some aspect of sleep homeostasis. Other than these early changes, SampEn remained stable throughout the 22-hour recording period in both wake (Fig. 8C) and NREM (Fig. 8D) at baseline and day 1 post-stroke in MCAO rats. Although SampEn showed evidence of circadian modulation, inter hemispheric MSE asymmetry (contralesional/ipsilesional SampEn) was not significantly affected by time of day at baseline or on D1 post-stroke (Fig. 8D, E).

**Figure 8:**
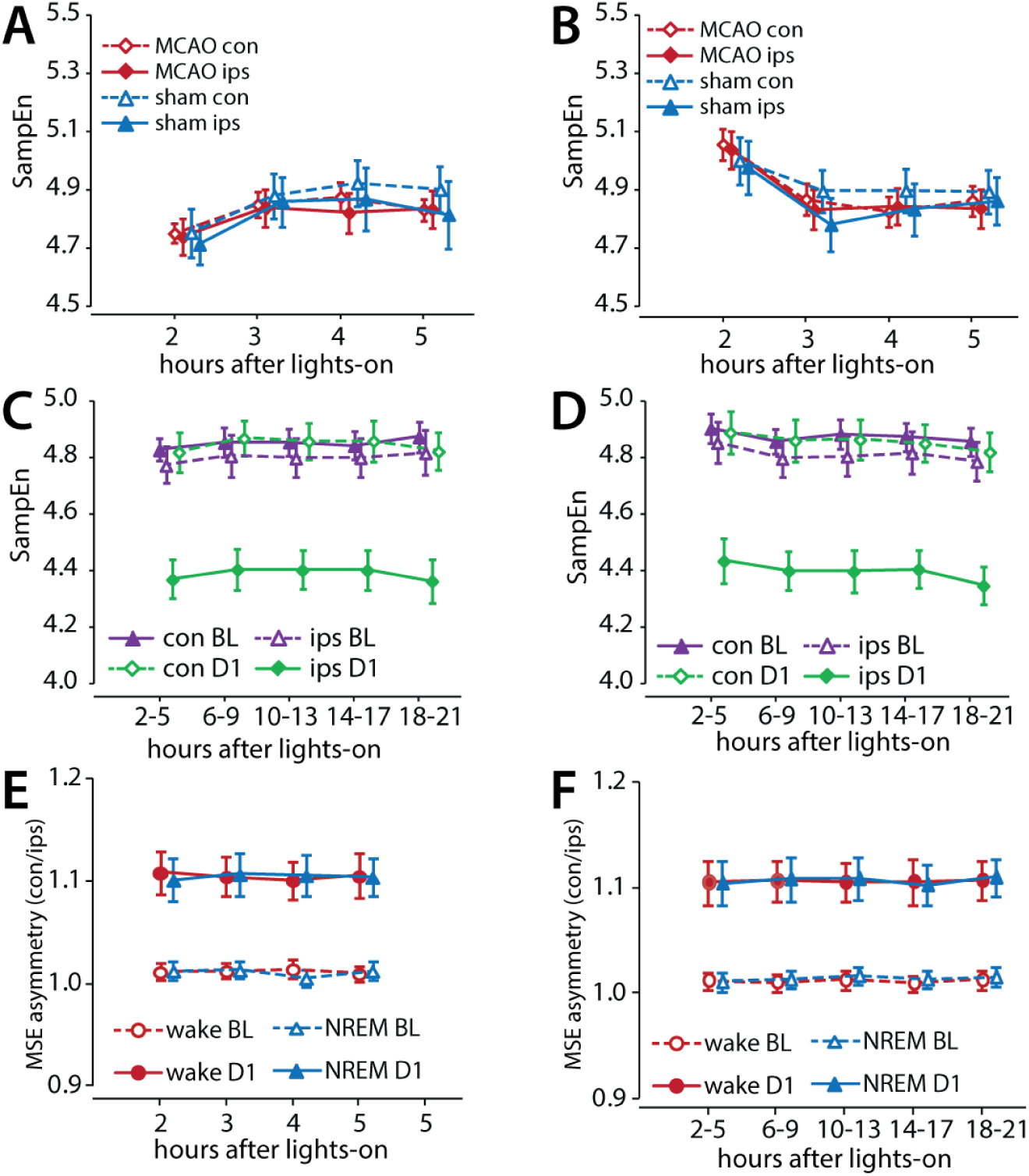
Circadian effects on SampEn and MSE asymmetry. **A:** Mean wake SampEn in the first 5 hours after lights-on in ipsilesional (ips) and contralesional (con) hemispheres of stroke (MCAO) and sham animals at baseline. Values are mean ± s.e.m. **B:** Mean NREM SampEn in the first 5 hours after lights-on in ipsilesional (ips) and contralesional (con) hemispheres of stroke (MCAO) and sham animals at baseline. Values are mean ± s.e.m. **C:** Mean wake SampEn in 4-hour bins at baseline (BL) and on D1 in MCAO animals. ips = ipsilesional, con = contralesional. Values are mean ± s.e.m. **D:** Mean wake SampEn in 4-hour bins at baseline (BL) and on D1 in MCAO animals. ips = ipsilesional, con = contralesional. Values are mean ± s.e.m. **E:** MSE asymmetry in wake and NREM (contralesional/ipsilesional SampEn) during the first 5 hours after lights-on in MCAO animals at baseline (BL) and on D1.

## Discussion

Many neurological disease states have been associated with reduced complexity, or reduced information content, in EEG signals [13–20,22,24,26]. Thus, loss of complexity seems to be an aspect of many pathological dynamics. Conversely, it seems likely that, the degree of complexity within a signal could be related to the ability of a system to recover from damage. EEG complexity could therefore be used as a diagnostic or predictive tool in clinical or pre-clinical settings and aid in the planning and implementation of optimal rehabilitation.

Complexity can be defined as meaningful structure or the information content within a signal [11]. While this intuitive definition is quite straightforward, there are many different approaches to quantifying complexity. Both Lempel-Ziv complexity analysis [10] and multiscale entropy analysis [11,12] detect thenumber of repeating sequences within a signal as a measure of regularity or predictability of a time series. Irregularity, however, does not necessarily imply complexity: a random signal such as white noise has by definition no meaningful information content, but is highly irregular and therefore has high Lempel-Ziv complexity. By contrast, information-rich signals such as the EEG don’t have very high LZC values. Sample entropy, as calculated via multiscale entropy analysis, aims to quantify complexity so that both completely predictable and completely random signals results in low complexity values [12], resulting in high values for EEG and lower values for white noise, as well as sine waves. As a result, sample entropy and Lempel-Ziv complexity of the same signal are not closely correlated. Thus, although these different measures of complexity are both assumed to reflect the information content of the same signal and to point to possibly identical underlying neuronal functions, the mechanistic implications of altered signal complexity remain somewhat nebulous.

In this paper, we compared the use of LZC and sample entropy analysis in quantifying changes in EEG complexity after stroke, as well as the use of these measures in predicting recovery. As in stroke patients [22,24,26,27], cortical strokes in rats led to reduced EEG complexity in the ipsilesional hemisphere, but not in the contralesional hemisphere. This was the case for both LZC and SampEn, although both complexity measures show different temporal dynamics. LZC was acutely reduced and returned to baseline levels 4 days after stroke. SampEn, by contrast, remained low much longer and returned to baseline and sham levels only after more than 10 days. As such, MSE could be used to identify the lesioned hemisphere in subchronic and chronic phases after stroke in rats as well as in patients [24].

Although SampEn was more severely affected by stroke than LZC, acute changes in SampEn were not correlated to functional recovery. By contrast, acute LZC asymmetry was correlated to the amount of recovery 31 days post-stroke in all vigilance states, providing a window for predicting behavioral outcome acutely after stroke. Likewise, LZC has been shown to outperform entropy measures such as SampEn in predicting behavioral function in other conditions [20,23]. Because LZC was only reduced acutely after stroke in rats and recovered quickly, it had little value as a predictor for behavioral outcome in later EEG recordings. However, because stroke recovery in rodent models is typically much faster than in humans [31], LZC may remain altered for a longer time after stroke in patients. If such a prolonged change is indeed present, this this provides an extended window for the use of LZC in predicting functional recovery in patients. If an aspect of the EEG is to be easily useable for predicting recovery, it is ideally not very sensitive to factors that are not related to altered brain or behavioral function. Unlike spectral power, inter-animal variability is quite low for both LZC and SampEn, even after stroke. A baseline measure or other normalization is not therefore not necessary to obtain interpretable LZC asymmetry values. Both measures are also independent of signal amplitude, which can be affected by factors such as age and sex [7–9]. However, both LZC and SampEn are not simply reflective of power spectral changes in the EEG.

Additionally, a useable predictor should remain relatively stable during a recording period. This seems contradictory to the greater sensitivity of complexity measures in detecting EEG dynamics compared to time-frequency analysis [21]. Both SampEn and LZC showed circadian changes in wakefulness and NREM sleep that are possibly related to sleep homeostatic processes. Although time-of-day effects on complexity were much smaller in LZC than in SampEn, sleep deprivation has previously been shown to affect LZC in NREM sleep [21]. Sleep disturbances are common in stroke patients [4], but did not occur in the animals in the current study [32]. However, unless largely asymmetrical wake activity occurred [33–35], sleep loss likely has similar effects on the EEG in the damaged and healthy hemisphere. In this study, fine motor performance, an asymmetric behavior, was tested on the day after EEG was recorded. As these skilled reaching sessions lasted fewer than 15 minutes and have been shown to have only have short-lasting effects on the sleep EEG [34], it is unlikely that this affected EEG complexity recorded on the following day. Moreover, wake LZC remained unaffected by sleep deprivation, even though variability between epochs increased [21]. Thus, resting wake EEG LZC asymmetry is likely stable even in the context of sleep-wake disturbances and may thus be the best choice for use in patients.

